# Ghost QTL and hotspots in experimental crosses - novel solution by mixed model with nonzero mean

**DOI:** 10.1101/825562

**Authors:** Piotr A. Szulc, Jonas Wallin, Małgorzata Bogdan, R.W. Doerge, David O. Siegmund

## Abstract

“Ghost-QTL” are the false discoveries in QTL mapping, that arise due to the “accumulation” of the polygenic effects, uniformly distributed over the genome. The locations on the chromosome which are strongly correlated with the summary polygenic effect depend on a specific sample correlation structure determined by the genotype at all loci. During the analysis of e-QTL data or recombinant inbred lines this correlation structure is preserved for all traits under consideration, and may lead to the so called “hot-spots” via the detection of the summary polygenic effect at exactly the same positions for most of the considered traits. We illustrate that the problem can be solved by the application of the extended mixed effect model, where the random effects are allowed to have a nonzero mean. We provide formulas for estimating the thresholds for the corresponding t-test statistics and use them in the stepwise selection strategy, which allows for a simultaneous detection of several QTL. Extensive simulation studies illustrate that our approach allows to eliminate ghost-QTL/false hot spot effects, while preserving a high power of detection of true QTL effects.

## 1 Introduction

It has been strongly suggested that many of the quantitative traits, including gene-expression levels, are subject to polygenic variation (Price *et al.* 2008; Fraser *et al.* 2010; Fraser *et al.* 2011; Turchin *et al.* 2012; VilhjÁ lmsson and Nordborg 2013). The classical infinitesimal genetic model, which assumes that polygenes are uniformly distributed over the whole genome, “forms the basis of quantitative genetics theories (Bulmer 1980; Henderson 1988; Falconer and Mackay 1996) that have been applied successfully to genetic improvement of livestock” (Liu and Dekkers 1998). In case of experimental crosses, where the parental lines are usually very much different with respect to many relevant features, it is natural to assume that the average polygenic effect may be different between these lines. As discussed in Deckers and Dentine (1991); Visscher and Haley (1996); and Liu and Dekkers (1998), the individually small effects of the polygenes can “accumulate” at certain positions on the chromosome, where they may lead to detection of ghost QTL. As noted by Visscher and Haley (1996) this effect can not be eliminated by conditioning on marker cofactors, as suggested by Composite Interval Mapping (Zeng 1994).

In this article we show that the ghost QTL positions depend mainly on the structure of the “design” matrix of genotypes. In case when many traits are regressed on the same genotype matrix, as in recombinant inbred lines or e-QTL mapping experiments, this may lead to hot-spot effects, i.e. the appearance of ghost QTL at the same positions for a large number of different traits. In this paper we show that ghost QTL can be eliminated by application of mixed effect model, where we allow the random effects to have a non-zero mean. The non-zero mean allows a polygenic influence on the difference in mean trait values between different inbred lines and to some extend plays the role of a fixed effect describing population assignments, often used to eliminate confounding in Genome-Wide Association Studies (see e.g. Yu *et al.* (2006); Zhao *et al.* (2007); VilhjÁ lmsson and Nordborg (2013)). We investigate the distribution of t-test statistics in this mixed effects model and demonstrate that the correlations between these statistics at neighboring genomic locations are substantially weaker than between the corresponding test statistics in the classical single marker modeling. However, similarly to the classical model, trajectories of the t-statistics can still be well approximated by the Ornstein-Uhlenbeck process. This allows us to calculate threshold values to control a weak sense Family Wise Error Rate at the nominal level. We use these critical values in a step-wise selection procedure, which allows for simultaneous detection of major QTL and has much better properties than the respective single marker tests. We use simulation to demonstrate that our method compares favorably to other approaches for eliminating confounding effects, for example, the classical mixed model with a zero mean or principal components of the design matrix as supplementary regressors. We show that our approach allows us to eliminate ghost QTL and yet yields high power to detect major QTL. Additionally, our method allows for a substantial improvement in precision of QTL localization as compared to the classical fixed effect model approach. We apply our method to the data of (Zeng *et al.* 2000) to reveal the genetic architecture of the shape of the posterior lobe of the male genital arch in *Drosophila*. According to our analysis the relatively large (72%) heritability of this trait can be attributed entirely to the polygenic effects and most (if not all) QTL identified in earlier articles can be considered as ghost QTL.

## 2 Methods

### 2.1 Statistical Model for QTL mapping with polygenic effects

The main purpose of our statistical analysis is the separation of a few large QTL from the polygenic background. We consider a backcross population, and we code the genotypes as *X*_*i j*_ = 1 if the *i*-th individual is homozygous at the *j*-th locus and *X*_*i j*_ = − 1 if it is heterozygous at this locus. Our model assumes that polygenic effects are uniformly distributed over the whole genome. Mathematically, it can be written

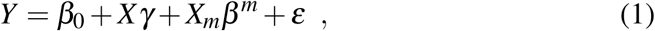

where *Y* is the *n* dimensional vector of trait values, *X* is the *n* × *p* matrix with genotypes of polygenic loci, *γ* is a *p*-dimensional vector from the multivariate normal distribution, *𝒩* (*µ***1**, *τ*^2^**I**_*p*×*p*_), *X*_*m*_ is the matrix containing genotypes associated with *m* large QTL effects, *β*^*m*^ ∈ *R*^*m*^ and *ε* ∼ *𝒩 (***0**, *σ* ^2^**I**_*n*×*n*_). The novelty of our model is that we allow *µ* to be different from zero, which permits the polygenes to affect the difference between expected values of the trait in the parental inbred lines.

The model (1) can be also written as

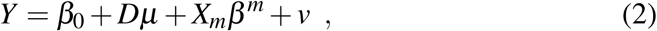

where 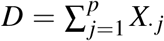 is the vector containing the sum of all columns of *X* and *v* ∼ *𝒩* (0, *σ* ^2^*I* + *τ*^2^*XX*^*T*^) is the random vector containing the random noise and the variance components of the polygenic effects.

### 2.2 Estimation and testing

The parameters *τ* and *σ* in the model (2) can be estimated using the method of the restricted maximum likelihood (REML). For this purpose we let Σ = *σ* ^2^*I* + *τ*^2^*XX*^*T*^. REML estimates parameters of the model (2) by maximizing the restricted likelihood, which up to an additive constant can be written as

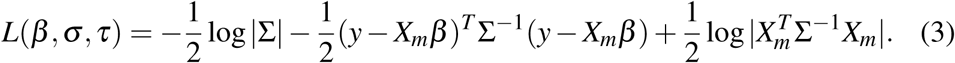

For fixed values of *τ* and *σ*, the value *β* maximizing the likelihood *L*(*β, σ, τ*) is given by

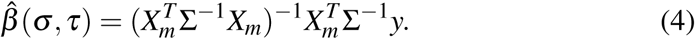

Replacing *β* with 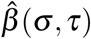 in (3) we obtain

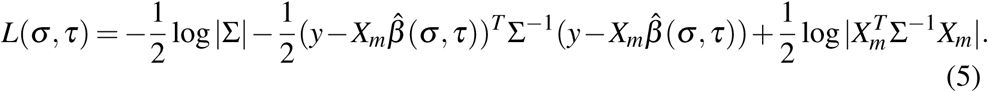

REML estimates 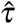 and 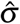 are the ones which minimize (5).

#### Remark 1

*The computational complexity of evaluating the log likelihood in* (5) *is 𝒪* (*n*^3^ + *m*^3^). *This can be quite prohibitive when we take into account that it has to be repeated a large number of times with different matrices X*_*m*_. *To improve the efficiency of the optimization one can compute the singular value decomposition of X* = *USV*^*T*^ *once, with computational complexity of 𝒪* (*n*^3^ ∧*p*^3^)*). Using the decomposition, the log likelihood can be evaluated as*

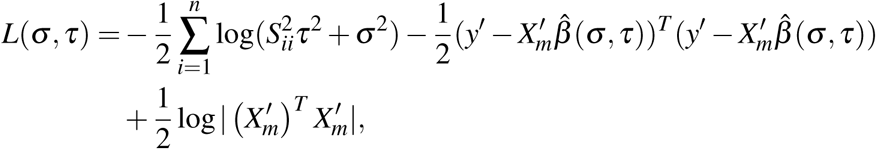

*where* 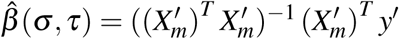, *and*

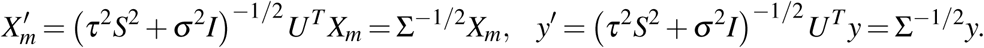

*Since S is a diagonal matrix the complexity of evaluating the log likelihood is now 𝒪* (*nm* + *m*^3^). *Further improvement of efficiency can be obtained by application of the Sherman-Morrison formula, which allows to reduce m*^3^ *to m*^2^. *We will not explore this techniquet here since m is typically small.*

Once the parameters are estimated we can perform the t-test at every locus in the following manner: We multiply the matrix *X*_*m*_ and the vector *Y* by 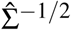:

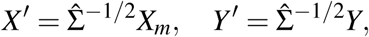

so we obtain

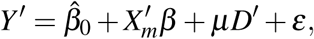

where 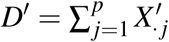 and *ε* ∼ *𝒩* (0, *I*). Now we can estimate the parameters *β*_0_, *β* and *µ* using a regular multiple regression model and calculate the *t*-test statistic to examine the significance of the *j*-th genetic locus included in the design matrix *X*_*m*_,

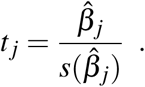

### 2.3 Step-wise selection procedure

Since we do not know which loci should be included in the matrix *X*_*m*_, we employ a modification of the step-wise selection procedure to identify large QTL. The procedure consists of two steps: forward selection and backward elimination. In the forward selection step we start with *X*_*m*_ = [1] (a column of ones) and estimate 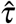 and 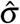. Then we perform the likelihood ratio test for the hypothesis *H*_0_ : *τ* = 0 and set 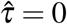 if this hypothesis is not rejected. Subsequently, we fit a sequence of *p* single locus models (2), where *X*_*m*_ is supplemented with just one genetic locus at a time, and we add to the model a marker with a highest value of the t-test statistic. Then we recompute 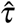 and 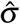 and repeat the search procedure looking for the genetic marker that would lead to the greatest improvement over the current model. We repeat this until the next locus to be added is not significant with respect to the Bonferroni correction at a liberal Family Wise Error Rate (FWER) of 0.25. Finally, the set of selected loci is filtered out using the backward elimination procedure, with the critical value adjusted to multiple testing according to the procedure described in the following section. After each of the backward elimination steps we re-estimate *τ* and *σ* and test if *τ* is significantly different from 0.

### 2.4 Multiple testing adjustments

Since the identification of important loci is based on an extensive search over the whole genome, the critical values for the respective test statistics need to be adjusted for multiple testing. Since the t-test statistics at neighboring loci are positively correlated, the Bonferroni correction is unnecessarily conservative. In the case of a backcross design the sequence of test statistics at consecutive locations can be approximated by an Ornstein-Uhlenbeck process (see e.g. [Feingold, Brown and Siegmund (1993), Dupuis and Siegmund (1999)] and [Siegmund and Yakir (2007)]), where the correlation between t-statistics at neighboring loci decays exponentially with the genetic distance between these loci: log*Cor*(*t, s*) = − *δ* |*t* − *s*|. In [Siegmund and Yakir (2007)] this approximation is used to calculate the critical value *t*_*crit*_ to control FWER at a level *α* by numerically solving the (approximate) equation

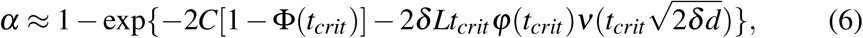

where Φ(·) and *φ*(·) denote the cumulative distribution function and density of the standard normal distribution, *C* is a number of chromosomes, *L* is a total genetic length (in cM), *d* is the average distance between neigboring loci (in cM) and

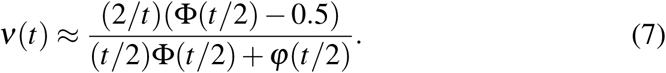

In case of the backcross under a regular fixed effects model, the coefficient *δ* can be calculated analytically and is equal to 0.02 (see e.g. [Dupuis and Siegmund (1999)]). In the presence of the polygenic effects (2), the intricate structure of dependencies between *t* statistics at neighboring loci substantially complicates the theoretical analysis of the correlation decay. Instead, we empirically verified that the decay is still roughly exponential, where the rate of correlation decay increases with *n* and the unknown variance of random effects *τ*^2^ (see Figure 1). In our mapping procedure at each step of the backward elimination we estimate *δ* based on the empirical decay of the correlations between neighboring test statistics and then calculate a significance threshold using the formula (6).

**Figure 1:**
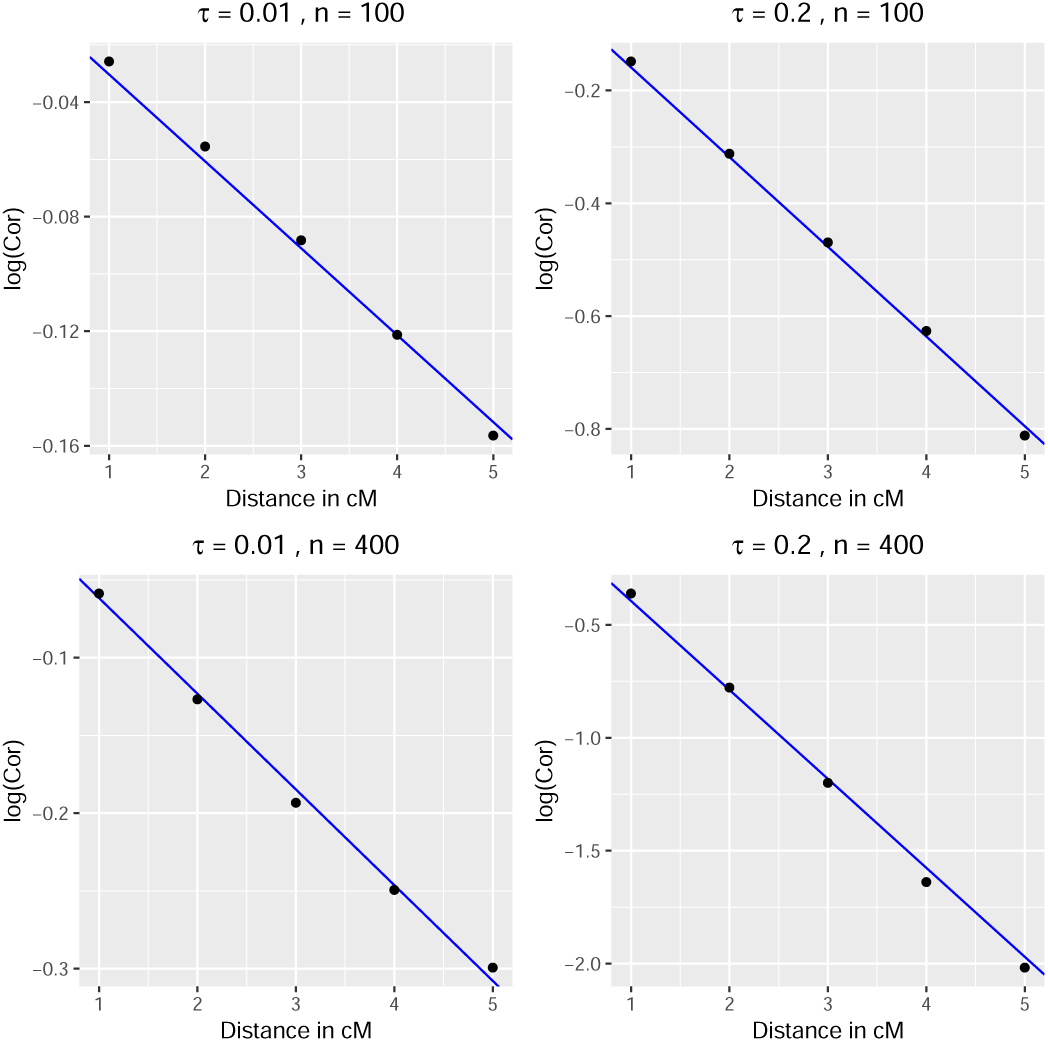
The distance between markers (in cM) and the logarithms of correlations between *t*-values. The calculations are based on known *τ* and *σ* = 1.

To verify that our procedure controls the probability of incorrectly including false QTL, we performed 1000 replicates of the experiment in which we simulated the data according to the model (2) with the vector *β* = 0. We simulated 10 chromosomes, each of length 150 cM, with polygenic effects placed uniformly every 1cM. The single marker test statistics were also calculated at markers spaced every 1cM. In Figure 2 we present the percentage of replications under which at least one of the single marker statistics in the mixed effect model was significant (Family Wise Error Rate, FWER). The figure shows that in an ideal case when *τ* and *σ* are known (i.e. we do not have to estimate them), FWER is very close to the nominal value of 0.05. When these parameters are estimated, FWER is slightly below that level when *τ* = 0 and slightly above when *τ* is large. The performance under *τ* = 0 results from the fact that in some small proportion of cases the null hypothesis that *τ* = 0 is rejected and the threshold is unnecessarily shifted up. In case when *τ* is large we observe the opposite effect, which suggests that *τ* is slightly underestimated. However, in both cases FWER is close to the nominal level and seems to converge to this level when *n* increases.

**Figure 2:**
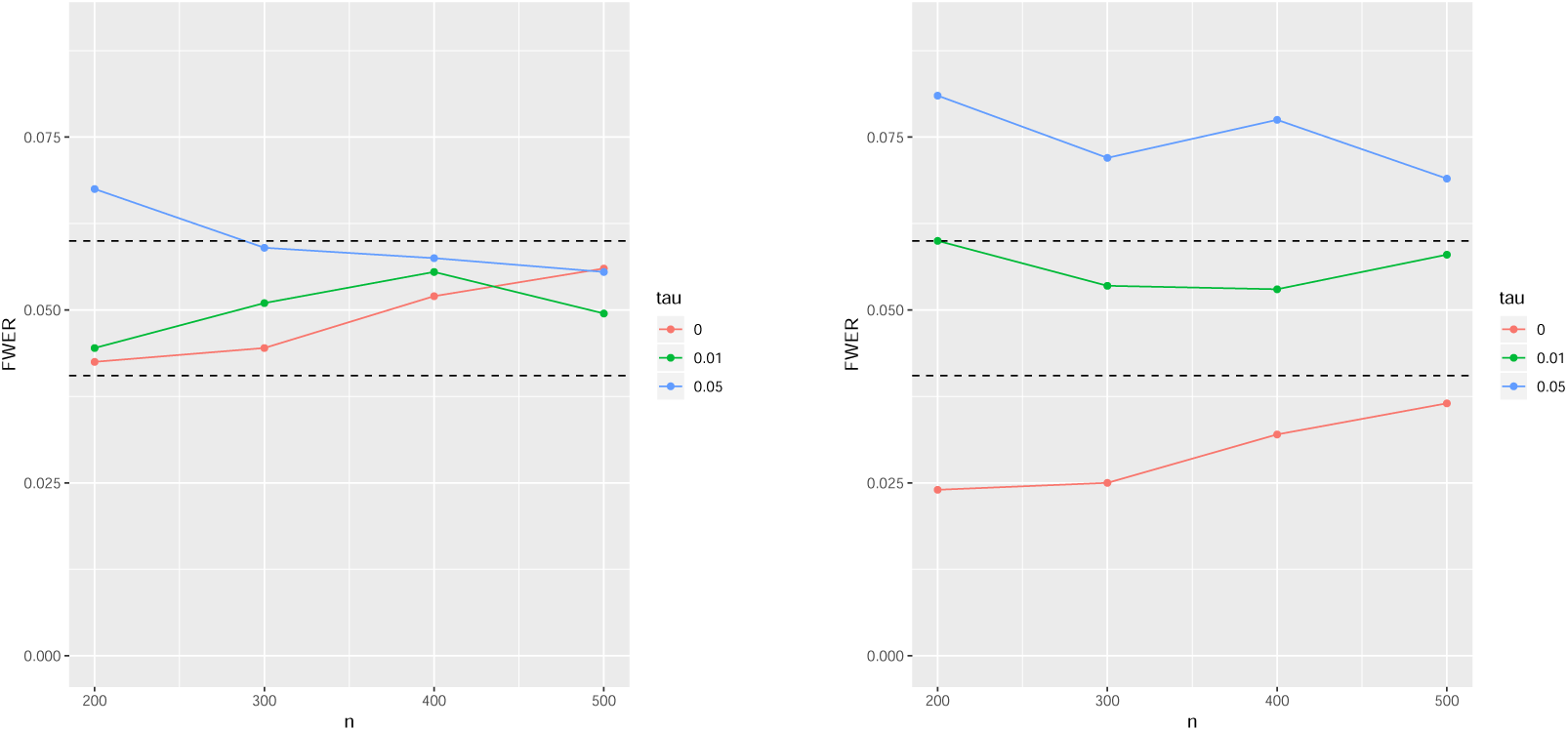
FWER for different *τ* and *n*. On the left we assume that *τ* and *σ* are known, on the right they are estimated. Horizontal bars represent the 95% error bands.

### 2.5 Markers vs polygenic loci

Clearly the markers used for QTL mapping do not need to coincide with polygenic loci. In our simulations we considered two scenarios. In both of them we simulated 10 chromosomes of the length of 150 cM, with polygenic loci spaced uniformly every 1cM. In the first set of simulations the genetic markers were placed exactly at the positions of these loci. In the second set of simulations we tried to capture the polygenic effects using markers uniformly spaced at the distance of 5cM.

### 2.6 Methods to compare

We compared our approach based on the mixed model with nonzero mean with several other methods often used in QTL mapping.

- **Single marker approach based on the regular fixed effects model.** Here we perform regular single marker t-tests adjusted for multiple testing using the threshold provided in [Dupuis and Siegmund (1999)].
- **Step-wise selection strategy based on the fixed effects model:**

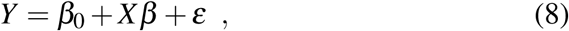

with *ε* ∼ *N*(0, *σ* ^2^*I*).
- **Step-wise selection strategy based on the mixed effects model with zero mean:**

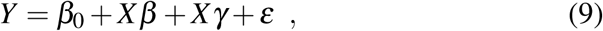

with *γ* ∼ *N*(0, *τ*^2^*I*) and *ε* ∼ *N*(0, *σ* ^2^*I*).
- **Step-wise selection strategy based on the fixed effects model and including several principal components of the matrix X:**

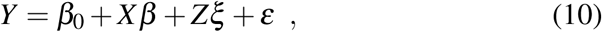

where *ε* ∼ *N*(0, *σ* ^2^*I*) and the columns of the matrix *Z* contain several first principal components of matrix *X*.
- **Step-wise selection strategy based on the mixed effects model with zero mean and including several principal components of the matrix X.**

### 2.7 Simulated genetic model

To compare statistical properties of different procedures we performed 1000 replicates of the experiment, where in each simulation we independently generated the backcross design matrix *X* and the trait values according to the model

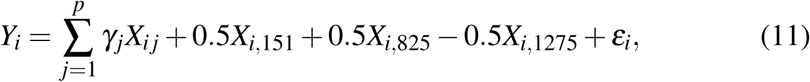

with *p* = 1500 polygenic loci uniformly spaced over ten 150 cM chromosomes, *γ*_1_, ⋯, *γ*_*p*_ independently sampled from *𝒩* (0.004, *τ* = 0.01) and the random error *ε*_*i*_ ∼ *N*(0, 1). Three QTL are placed in positions 151, 825 and 1275, so the first one is at the beginning of a chromosome and two others in the middle of chromosomes. We considered two sample sizes, *n* = 200 and *n* = 400.

We summarize the results by providing the following characteristics

- Power of identifying each of the three QTL,
- FP -the average number of false positives,
- Average distance between identified locus and the true QTL position,
- Average value of 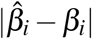.

Due to the strong correlation between neighboring loci, the precision of QTL localization is rather limited. Therefore, when calculating the power we consider a true QTL to be properly identified if the procedure selects the marker within the distance of 15 cM from its position. Every discovery which is not within a 15 cM distance from the true QTL is treated as a false positive.

### 2.8 Simulating polygenic gene expressions

Many e-QTL and Genome-Wide Association studies (GWAS) report the existence of the so called “hot-spots”, i.e. expression QTL or single polymorphisms that are associated with widespread changes in the expression of distant genes (Schadt *et al.* 2003; Schadt *et al.* 2008; Wu *et al.* 2008; Yvert *et al.* 2003; Breitling *et al.* 2008). While some of these “hot-spots” have been validated biologically, others are often not consistent and elusive (PÉ rez-Encisco 2004; de Koning and Haley 2005; Breitling *et al.* 2008). Several reasons for possible false detections of “hot-spots” are discussed in Leek and Storey (2007) or Breitling *et al.* (2008); some believe that the “hot-spot” phenomena are artifacts of the correlation between e-traits caused by factors which are not accounted for in the statistical models used for detection of SNP-trait association. Here we perform a simulation study that illustrates that “hot-spots” arise because of the polygenic background.

We use model (1) to simulate polygenic gene expressions, corresponding to all *p* = 1500 genetic loci spaced uniformly on 10 chromosomes of the length of 150 cM for *n* = 200 individuals. Here the genotype matrix *X* is the same for each of the expressions, while the polygenic effects for the *k*^*th*^ trait *γ* _*jk*_ are independently simulated from the normal distribution *𝒩* (*µ*_*k*_, *τ*^2^), with *τ* = 0.01. The expected values of the polygenic effects *µ*_*k*_, *k* = 1, ⋯, 1500, are simulated as independent random variables from the normal distribution *𝒩* (0, 0.007^2^). To model cis-effects we additionally generated “large” coefficients *β*_*kk*_ from the normal distribution *N*(0.5, 0.1^2^).

### 2.9 Real Data Analysis

We used the mixed model with nonzero mean to analyze the well known *Drosophila mauritiana* data of [Zeng *et al.*(2000)]. The purpose of this analysis was the identification of QTL influencing the shape of the posterior lobe of the male genital arch in *Drosophila*. These data were extensively analyzed in [Zeng *et al.*(2000)] and [Bogdan *et al.* 2008] using different approaches based on the fixed effects multiple regression models. These analyses report 17 QTL, roughly uniformly distributed over these two chromosomes, with two strongest QTL located close to the centers of these chromosomes. As reported in [Zeng *et al.*(2000)], the results of the analyses with the multiple regression models substantially differ from the results of the Composite Interval Mapping of [Zeng (1994)]. Moreover, [Bogdan *et al.* 2008] shows that the likelihoods of different multiple regression models are comparable and the positions of identified QTL differ substantially, depending on the number of assumed effects in the model.

In this article we report the analysis of this data set with the regular single marker tests, the Composite Interval Mapping of [Zeng (1994)] with a window of 20 cM and the step-wise selection procedure based on the mixed model (1) proposed in this manuscript. The data includes genotypes of 39 markers on 2 autosomes for *n* = 491 individuals. Similary as in [Bogdan *et al.* 2008] we perform our analysis using *m* = 161 pseudo-marker explanatory variables spaced every 2 cM. Values of these pseudo-markers are calculated as the conditional expectations of the corresponding genotypes given the genotypes of observed flanking markers, as in the regression interval mapping of [Haley and Knott (1992)].

## 3 Results

### 3.1 Graphical comparison of different procedures

In Figure 3 we can see the graphical comparison of different methods when applied to the analysis of the trait simulated according to the model (1). In the upper left panel we can see that the regular single marker test analysis can identify QTL1 and QTL2 but misses QTL3, whose effect is opposite to the summary polygenic effect. This analysis generates also two significant “ghost” QTL at chromosomes 1 and 10. Moreover, the upper middle panel shows that the number of ghost QTL increases to five (pink vertical lines) when the data are analyzed with the stepwise selection procedure based on the fixed effects model. This phenomenon is due to the reduction of the variance of the residual error obtained when replacing the simple regression model with the multiple regression. In the middle left panel we can observe that the single marker tests within the mixed effects model (1) with *µ* fixed at 0 can identify only QTL2 but they also do not detect any false QTL. After reducing the residual variance by the stepwise selection strategy the mixed effects model with *µ* = 0 detects also QTL1 but still misses QTL3 and generates the ghost QTL at chromosome 1. The pink vertical curves in the lowest panel show that the stepwise selection strategy based on our proposed model (1) with *µ* ≠ 0 allows to identify all three QTL and does not generate any false QTL. The lower panel shows also the importance of using a proper selection strategy. On the left graph we can see that the single marker test analysis within the model (1) misses QTL1, while the middle graph shows that conditioning only on QTL2 and QTL3 generates the ghost QTL and the shoulder of QTL1. The right graph shows that this ghost QTL disappears when conditioning on all three QTL, properly identified by our stepwise procedure. Comparing upper, middle and lower panels we can also see that the mixed effects models allow for a higher precision of QTL localization as compared to the fixed effects model (narrower peaks on the trajectories of t-statistics).

**Figure 3:**
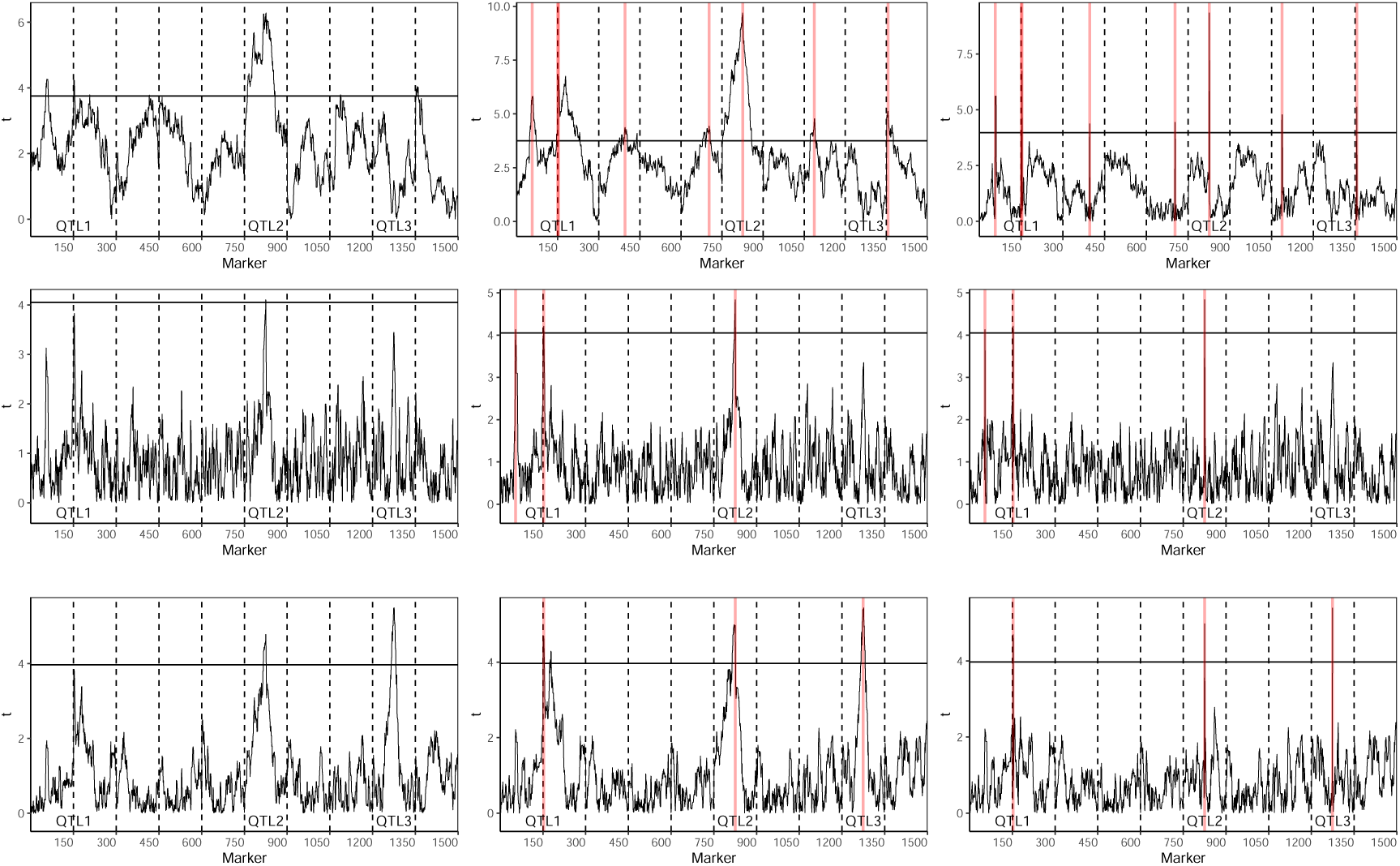
Results of QTL mapping for the trait simulated according to the model (11) with *n* = 200. The upper, middle and lower panels show the results of the analysis with the classical fixed effect model, the mixed model with *µ* = 0 and the mixed model (1), respectively. In each of these panels the graph on the left represents the results of the single marker tests, while the remaining panels present the results of the respective stepwise selection procedures. The pink vertical lines mark the positions of markers selected by these procedures. The middle graphs show the plots of t-statistics when conditioning on QTL identified on other chromosomes. The right graphs show the plots of t-test statistics conditioned on all identified QTL.

### 3.2 Power and the number of false positives

Table 1 provides the results of the comparison between different methods of QTL mapping for the data simulated according to the model (11). We provide the power of identification of each QTL, the average number of false positives (*FP*) and the false discovery rate (FDR). Moreover, in brackets we provide the average difference between the estimated and the true QTL position and the average value of 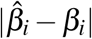.

**Table 1:**
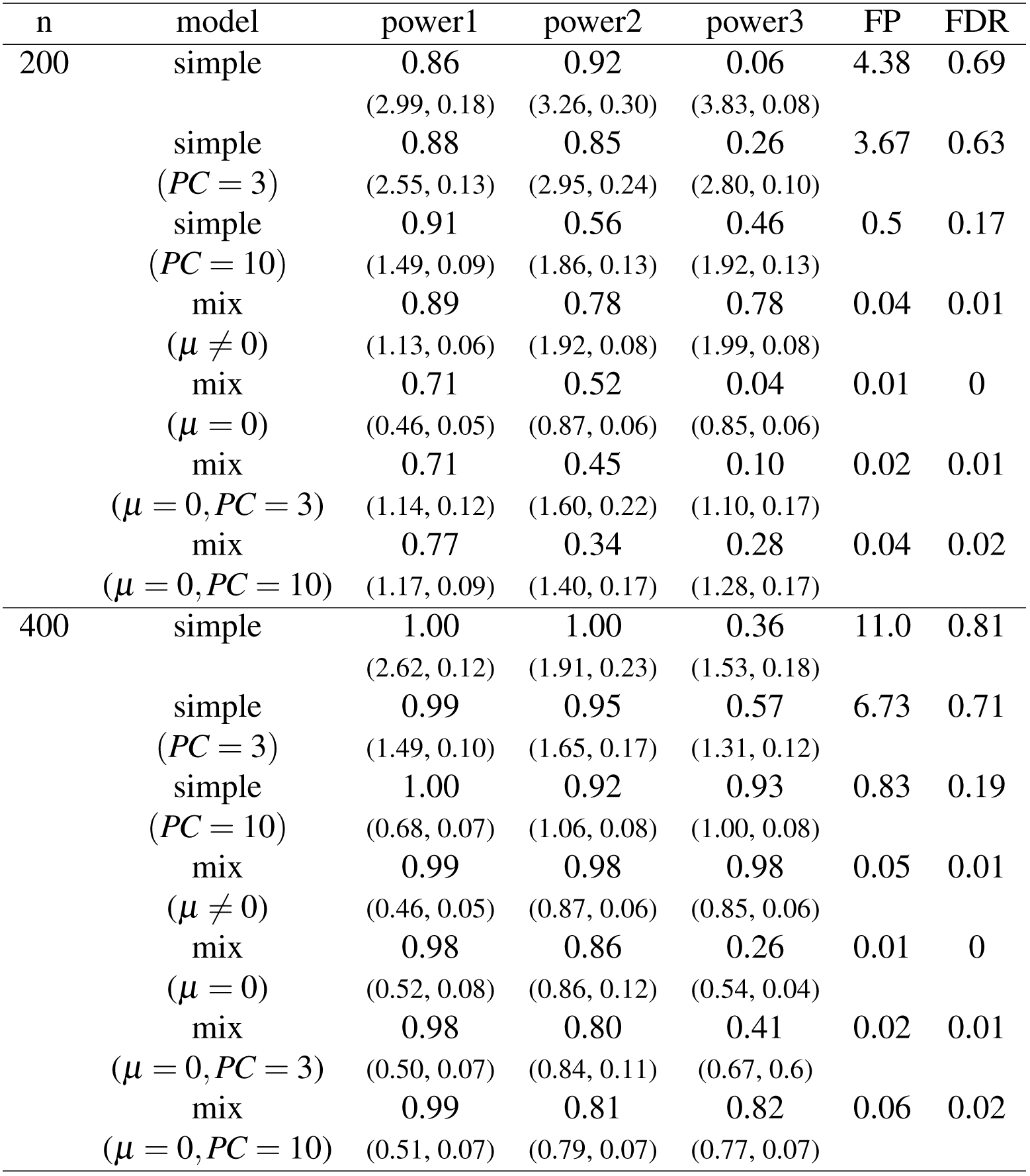
Statistical properties of different methods as applied to the analysis of data generated according to the model (11). We report Power, FP and FDR. Additionally, in the brackets we report a mean distance to real QTL and a mean value of 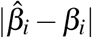.

Table 1 illustrates that indeed the model selection procedure based on the regular multiple regression model yields a large number of “ghost” QTL and that this number increases with the sample size. Moreover, this procedure is hardly capable of identifying QTL3, whose effect is of the opposite sign to the summary effect of polygenes. The number of “ghost” QTL is reduced when including several Principle Components (PCs) of the *X* matrix as covariates. However, to reduce FDR below 0.2 one needs to include 10 PCs, which for *n* = 200 leads to a substantial decrease of power of identification of QTL2. All approaches based on the mixed model allow to eliminate almost all “ghost” QTL. However, methods based on the models which assume that *µ* = 0 have a substantially smaller power than the procedure based on the model (1), with *µ* ≠ 0. Specifically, all methods which assume that *µ* = 0 have a rather small power of identification of QTL3. Similarly as in the case of the fixed effects models, adding PCs to the mixed model with *µ* = 0 leads to the reduction of the power of identification of QTL2 and improves the power of identification of QTL3. Here it is interesting to observe that the performance of both methods which use 10 PCs substantially improves with the sample size. The method based on the mixed model offers a better control over the number of false positives, which however comes at the price of some power loss.

It is also interesting to observe that the mixed effects models offer a substantially higher precision of QTL localization as compared to the classical fixed effects models. The precision of these classical models can be improved by including several first PCs of *X*. However, even then, the estimated locations are less precise then the ones provided by the mixed model approach (particularly for *n* = 400). The best precision of QTL location is obtained for the mixed model with *µ* = 0, which however comes at a prize of a substantial reduction of power as compared to the analysis with the model allowing for *µ* ≠ 0. Also, for *n* = 400, the mixed model with *µ* ≠ 0 provides a substantially better precision of estimation of QTL effects.

Table 2 provides results of the analysis when the polygenic effects are spaced every 1 cM but the search for QTL is performed using markers spaced every 5cM. In this case we observe that procedures based on the mixed model with *μ* = 0 suffer from the loss of power and loss of precision of the estimation of QTL effects. Instead, the procedure based on the mixed model with *μ* ≠ 0 is not affected by using sparsely spaced markers, which suggests that uniformly spaced markers can efficiently capture the mean of the polygenic effects even when the distance between these markers is rather large.

**Table 2:**
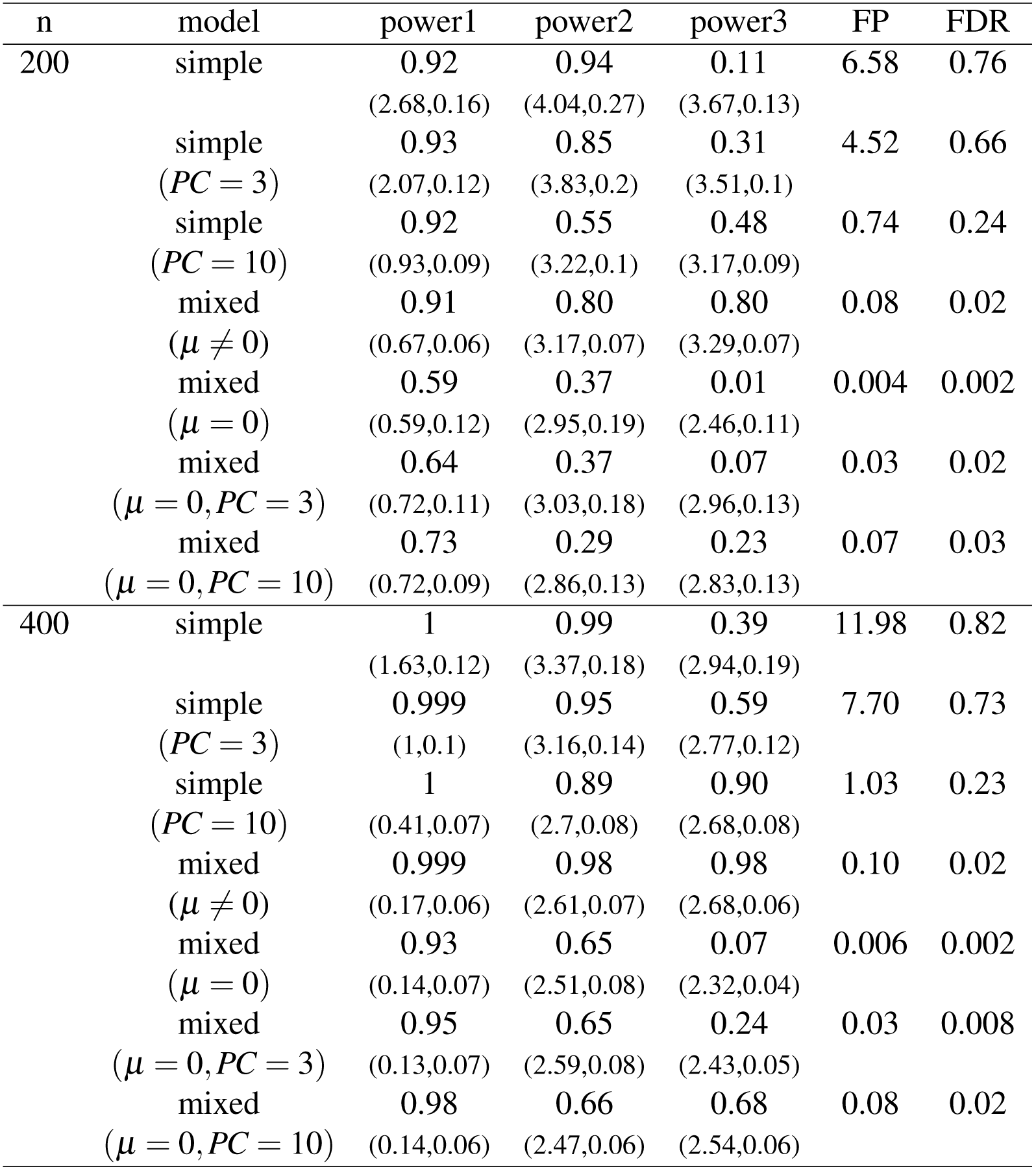
Statistical properties of different methods as applied to the analysis of data generated according to the model (11). We report Power, FP and FDR. In the brackets we report a mean distance to real QTL and a mean value of 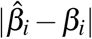. The polygenic effects are uniformly distributed at the distance of 1 cM but the search is performed over markers spaced 5CM.

Table 3 contains results of the analysis of the trait which is influenced only by few moderately sized QTL and has no polygenic background:

**Table 3:**
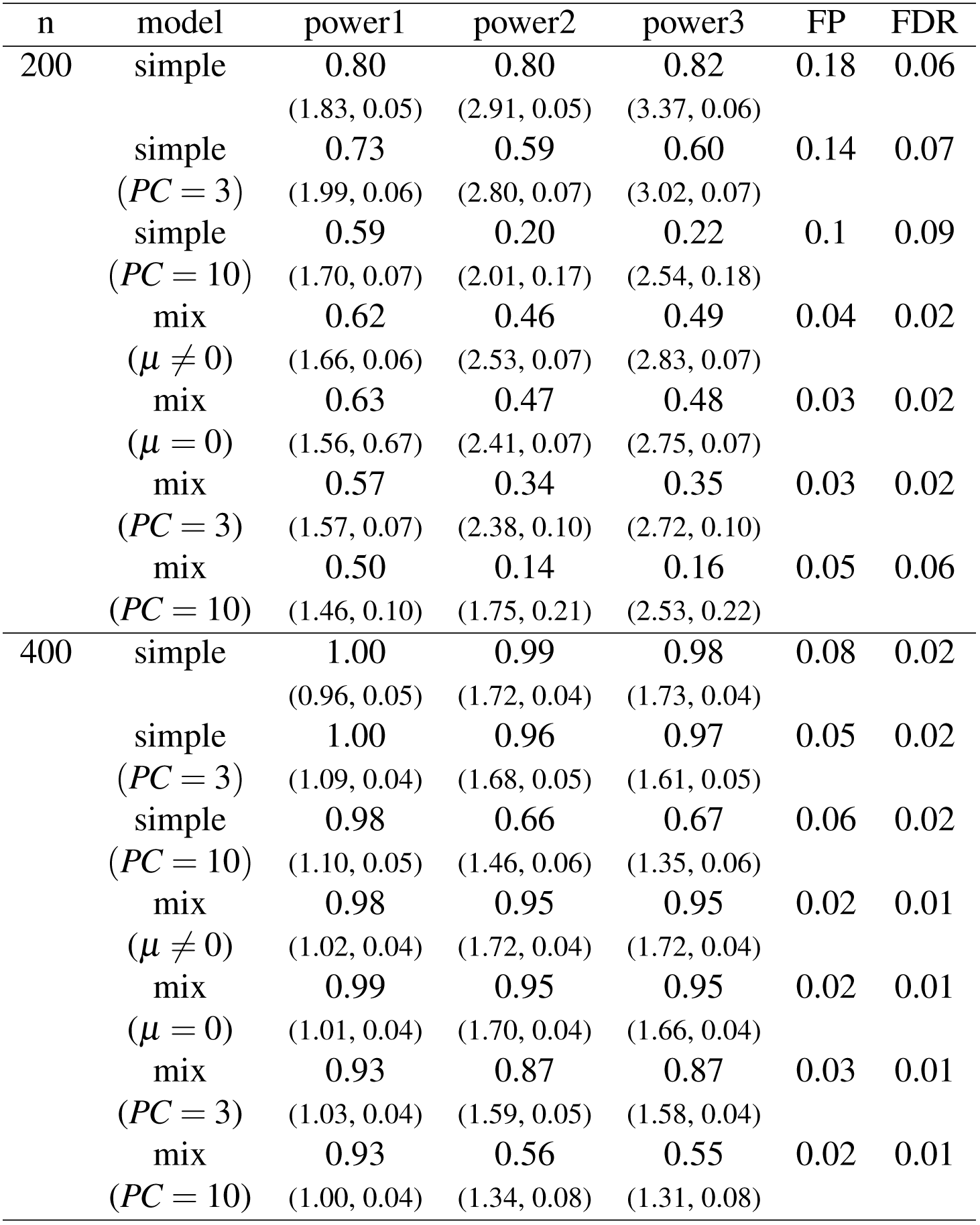
Statistical properties of different methods as applied to the analysis of the trait generated according to the model (12), with no polygenic background. We report Power, FP and FDR. In the brackets we report a mean distance to real QTL and a mean value of 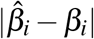.

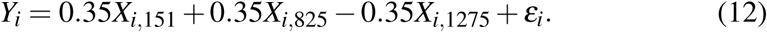

Here, we can see that the analysis based on the proper fixed effects model allows to obtain a highest power of QTL detection. In case of *n* = 200 we observe a substantial loss of power when the data are analyzed by the mixed model or/and when the model is supplemented by PCs. Interestingly, in this situation we also observe a substantially better precision of estimation of QTL location by the mixed model. The imprecision of localization of weak genetic effects by the fixed effects model is the reason for the slightly inflated number of false positives. This is because it sometimes occurs that the distance between the estimated and the true QTL location exceeds 15 cM, which is the threshold we use to classify true and false positives. When *n* = 400 both methods based on the mixed model compare very well with the fixed effects model. They yield only a slightly lower power and a slightly lower number of false positives. However, the deteriorating effect of including unnecessary PCs remains quite strong even for *n* = 400.

### 3.3 Hot-spots in e-QTL studies

In Figure 4 we see the results of the simulated e-QTL analysis, where for each of 1500 genetic loci we simulated polygenic expression levels according to the procedure described in Section 2.8. Red points mark the positions with coordinates *i* and *j* such that the single marker test for *j*^*th*^ expression and *i*^*th*^ locus is significant when using the multiple testing correction (6) at the Family Wise Error Rate *α* = 0.05*/*1500, adjusted to the number of traits by the Bonferroni correction.

**Figure 4:**
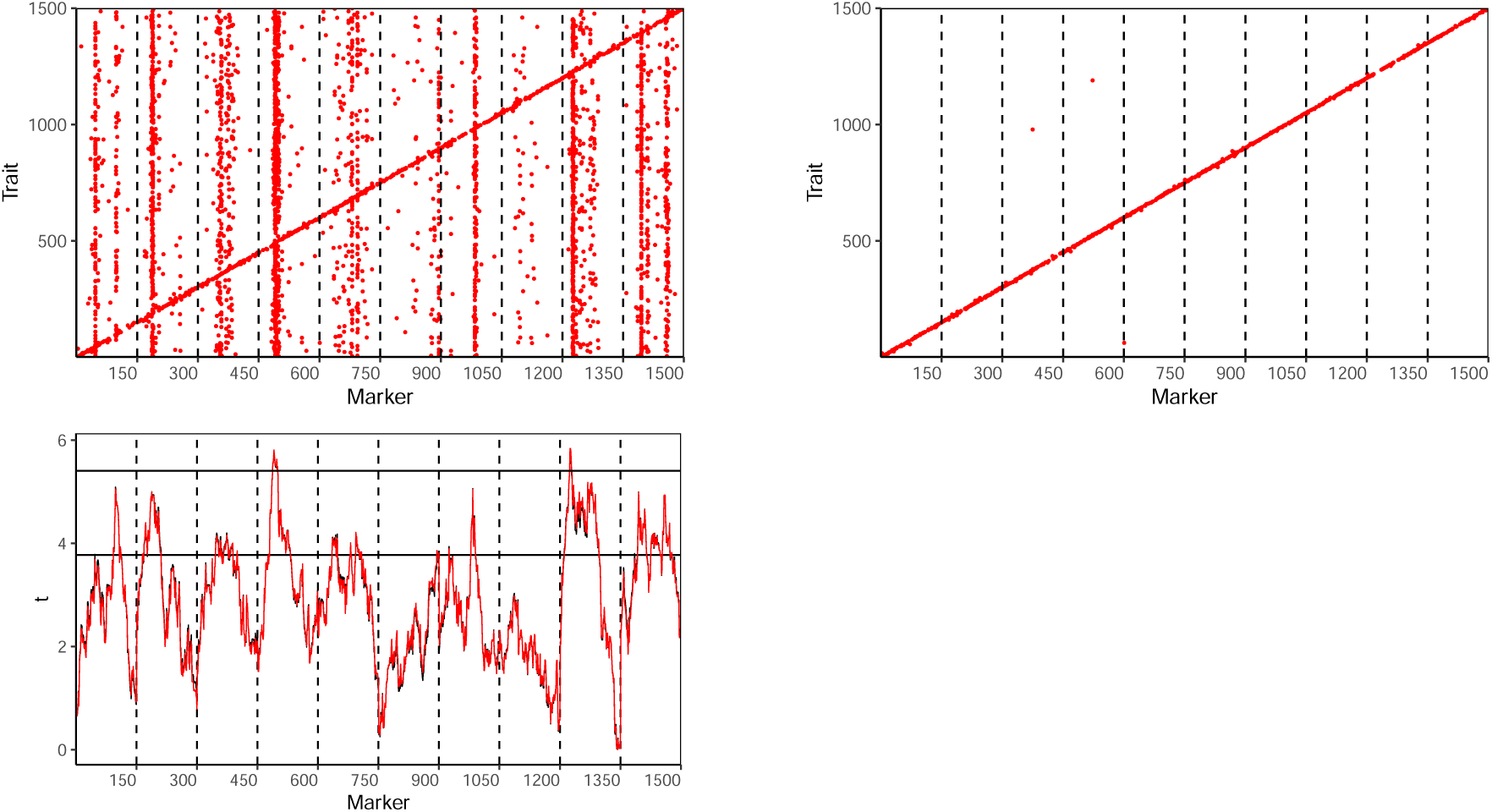
The upper panel represents the results of the analysis of the simulated e-QTL data with the classical single marker tests (left plot) and the step-wise selection strategy based on the model (1) (right plot). Values on the y-axis and x-axis correspond to the indices of the e-traits and the markers, respectively. Red points mark the positions which are significant at the level 0.05 after adjusting to the number of markers and the number of traits. In the lower panel we present means of the absolute values of *t*-test statistics over all 1500 traits (black line) and the theoretical expected values calculated according to the formula (14) (red line). In most positions these curves coincide completely. The horizontal lines correspond to the thresholds of [Dupuis and Siegmund (1999)] for FWER control at the levels 0.05 and 0.05*/*1500. The “upper” threshold uses Bonferroni correction to adjust to the number of traits.

The upper left panel represents the results of the analysis with the classical single marker tests. Here we can clearly see the diagonal line corresponding to cis-effects, but we also see ghost hot-spots, i.e. the vertical lines e.g. at the second, fourth and ninth chromosomes. These ghost hot-spots appear due to the fact that the design matrix *X*_*n*×*p*_ for e-QTL mapping is the same for for all gene-expression levels, which causes ghost QTL to appear in the similar positions for all these traits. Actually, given the design matrix *X* and the distribution of the polygenic effects one can approximate the expected values of the t-test statistics under the polygenic model of inheritance (see the formula (14) in the Appendix). In the lower panel in Figure 4 we show that these theoretical expected values perfectly agree with the mean of the absolute values of *t*-test statistics over all traits. We can also observe that these expected values exceed the multiple testing adjusted threshold at chromosomes 4 and 9, which suggests that e-QTL would be wrongly identified here for more than 50% of traits. This is confirmed by strong “hot-spots” at these positions, clearly visible in the upper left panel. Also, other visible hotspots at chromosomes 2, 7 and 10 agree well with peaks at the graph of expected values. The upper right panel illustrates that all ghost hot-spots are completely eliminated when analyzing the same data set with the step-wise procedure based on the mixture model (1).

### 3.4 Real data

Figure 5 contains results of the analysis of the Drosophila data of [Zeng *et al.*(2000)] with different methods of QTL mapping. In the left panel we observe that the absolute values of the single marker t-statistics exceed the critical value systematically over both chromosomes. These results suggest a strong polygenic background of the analyzed trait. In the middle plot we can see that the Composite Interval Mapping suggests a strong QTL close to the left flank of chromosome 1 and four or five suggestive QTL, roughly uniformly distributed over the chromosome 2. Other results of the analysis of this data set are reported in [Zeng *et al.*(2000)] and [Bogdan *et al.* 2008], which use a multiple regression models and suggest 17 QTL, roughly uniformly distributed over these two chromosomes. The right panel illustrates the results of the analysis of this data set with the mixed model (1), which attributes the whole 72% heritability of this trait to the polygenic background. We believe that this is a reasonable way of summarizing this data, taking into account lack of replicability of identified QTL positions by different methods based on the fixed effects model. It seems that precise identification of locations of so many QTL is here practically impossible, due to a limited sample size and a strong correlation between genotypes at neighboring loci.

**Figure 5:**
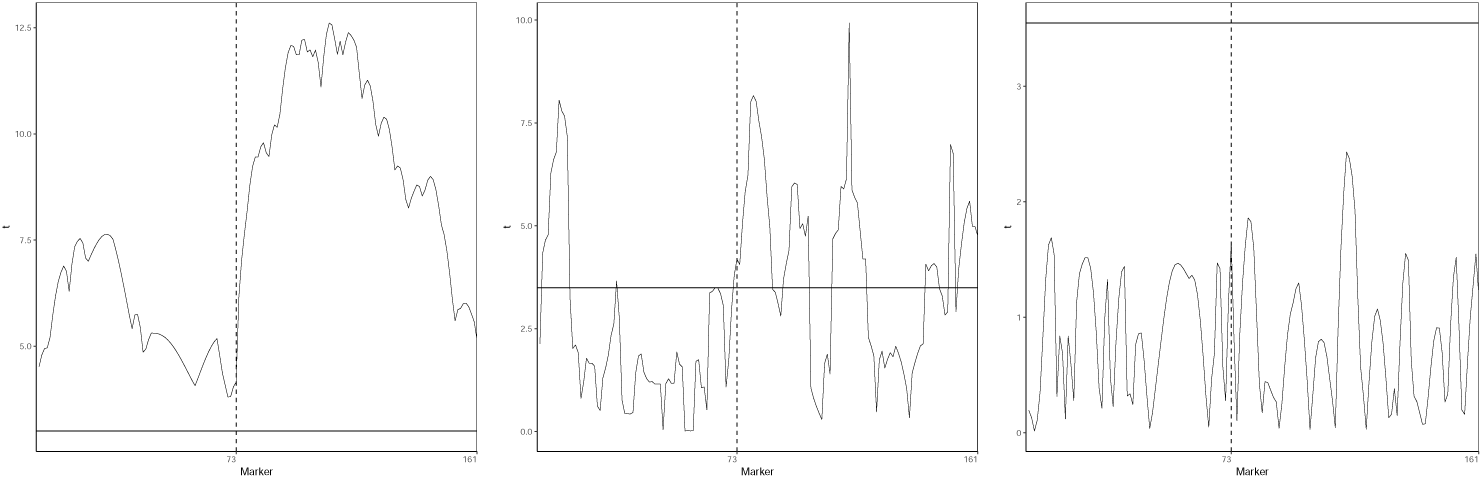
Results of the analysis of the *Drosophila* data of [Zeng *et al.*(2000)] with different methods of QTL mapping. Left panel represents single marker t-tests. The middle panel provides the results of the Composite Interval Mapping of [Zeng (1994)], where the t-statistics are calculated after conditioning over all markers out of the ± 20 cM window. The right panel represents the results of the single marker tests within the mixed model (1).

## 4 Discussion

It appears that “ghost-QTL” and the “hot-spots” often found in e-QTL analyses and GWAS may naturally arise in situations where the traits are influenced by a large number of polygenes, with marginally small individual effects. The complexity of inheritance of quantitative traits is far from a new concept, and has been well documented in both the animal and plant breeding/genetics literature. It has also been strongly suggested that many of the quantitative traits, including gene-expression levels, are subject to the polygenic adaptation (Price *et al.* 2008; Fraser *et al.* 2010; Fraser *et al.* 2011; Turchin *et al.* 2012), which forms the basis for the models used in our simulation study. The polygenic inheritance of the expression levels and be explained by the fact that the expression of a given gene is usually activated by the transcription factors produced by other genes. Thus, it depends not only on the genotype of its own cis-regulatory elements but also on the genotypes of cis-regulatory elements of the multitude of other genes involved in a genetic pathway.

With respect to “hot-spots”, it is our opinion that if viewed in the context of complex inheritance, they might be better understood and actually considered as false QTL that gather the effect of polygenes via the sample correlation matrix between genotypes of all loci. In the case of an e-QTL study, since the matrix of genotypes is the same for all e-traits such spurious associations tend to appear at the same location for different traits. Here, we have assumed that the polygenes are uniformly distributed over a single chromosome, and that their effects are independent random variables from the same normal distribution. Obviously, different patterns for the distribution of polygenes are possible, and have potential to lead to different patterns of “hot-spots”. As seen here, in the inbred lines the hot-spots resulting from the single marker/interval mapping analysis have a tendency to appear in the middle of the interval in which most of the polygenic effects have the same sign; and these hot-spots are relatively robust to the selection of different samples from a given experimental population. Comparing different experimental populations one expects higher reproducibility of hot-spots for recombinant inbred lines, where the marker genotypes in different lines are fixed and lead to the same “design matrix” for many experiments.

In this article we show that ghost-QTL and ghost hot-spots can be eliminated by the application of the mixed models with the non-zero mean of the random effects. The model used in this paper assumes that the intensity of the polygenic effects remains constant over the whole genome. This methodology can be naturally extended by allowing *μ* and *τ* to be smooth functions of location, which can be estimated for example by using the latent Gaussian process. We consider this as an interesting topic for a further research. Also, our simulations suggest that for large sample sizes the non-zero mean effect can be too some extend captured by inclusion of the relatively large number of Principle Components of the design matrix. It would be of some interest to quantitatively describe this phenomenon, which we leave as an interesting topic for a further research.

## Acknowledgments

We would like to thank Florian Frommlet for help in programming the inbred population simulation study, and Matthew Stephens for helpful remarks and references.

## 5 Appendix

### 5.1 Expected values of the single marker test statistics

Here we provide mathematical formulas for approximating the expected value of the single marker t-test statistics when the trait is influenced by polygenes according to the model (1).

First, recall that the least squares estimator of the regression coefficient in the simple regression model is

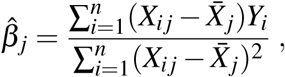

where 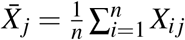.

Using the assumption that *β*_1_, …, *β*_*p*_ are independent random variables from the normal *𝒩* (*μ, τ* ^2^) distribution, we can write

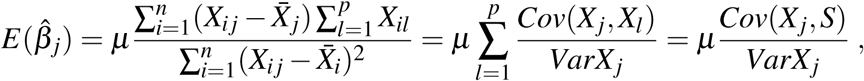

where *Cov*(*X*_*j*_, *X*_*l*_) is the sample covariance between the genotypes of *j*^*th*^ and *l*^*th*^ markers, *VarX*_*j*_ is the sample variance of the genotypes of *j*^*th*^ marker and *S* = (*S*_1_, …, *S*_*n*_) with 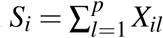.

Furthermore,

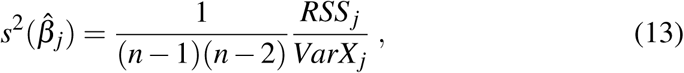

where *RSS*_*j*_ is the residual sum of squares in the simple regression model. Relying on Cochran’s theorem (see Section 1.3 of Shao 1998) it can be shown that given the vector of true regression coefficients *β, RSS*_*j*_ has a noncentral chi-square distribution *RSS* ∼ *𝒳*^2^(*n*− 2, *δ*_*d*_), with the noncentrality parameter

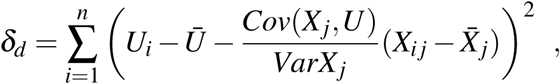

where 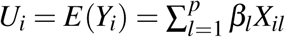 and *U* = (*U*_1_, …, *U*_*n*_).

Thus

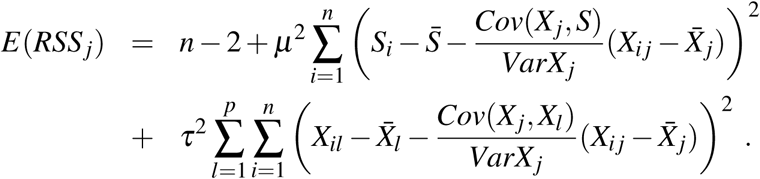

Subsequently, 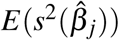 can be obtained by plugging *E*(*RSS*_*j*_) into the equation (13). To predict the hotspots for applications of a single marker test, or interval mapping, we rely on the fact that the expected value of the t-test statistics for the single marker tests can be well approximated by

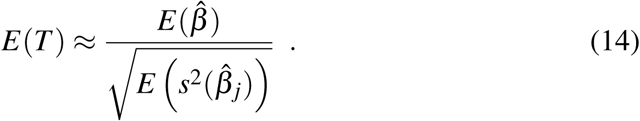

In Figure 4 the value *μ*^2^ was replaced by *E*(*μ*^2^) = 0.007^2^.

### 5.2 Heritability of the mixed model with non-zero mean

Let us define heritability as

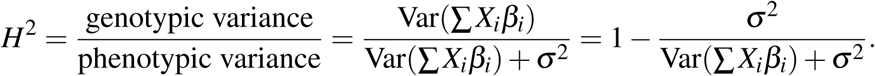

We have

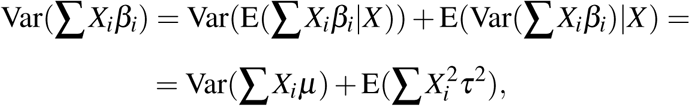

because *β*_*i*_ are independent and *β*_*i*_ ∼ *𝒩* (*μ, τ*^2^). If we have *c* chromosomes and each of them has *m* markers then we can write

- 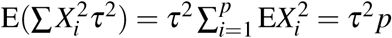 if *X*_*i*_∈{−1,1},
- 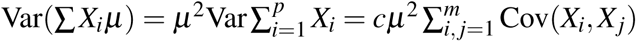

The calculations can be completed by observing that

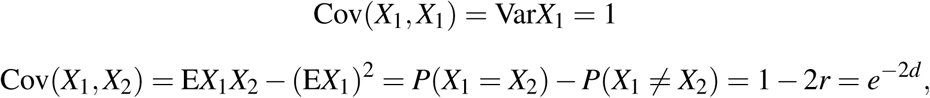

where *r* is the recombination rate and where *d* is a distance in Morgans. Similarly, Cov(*X*_1_, *X*_3_) = *e*^*−*4*d*^ and in general, Cov(*X*_*i*_, *X*_*j*_) = *e*^*−*2*d|i− j|*^.

